# Conformation of HIV-1 Envelope governs rhesus CD4 usage and simian-human immunodeficiency virus replication

**DOI:** 10.1101/2021.08.25.457697

**Authors:** Geraldine Vilmen, Anna C. Smith, Hector Cervera Benet, Rajni Kant Shukla, Ross C. Larue, Alon Herschhorn, Amit Sharma

## Abstract

Infection of rhesus macaques with simian-human immunodeficiency viruses (SHIVs) is the preferred model system for vaccine development because SHIVs encode HIV-1 envelope glycoproteins (Env) – a key target of HIV-1 neutralizing antibodies. Since the goal of vaccines is to prevent new infections, SHIVs encoding circulating HIV-1 Env are desired as challenge viruses. Development of such biologically relevant SHIVs has been challenging as they fail to infect rhesus macaques, mainly because most circulating HIV-1 Env do not use rhesus CD4 (rhCD4) receptor for viral entry. Most primary HIV-1 Env exist in a closed conformation and occasionally transit to downstream, open conformation through an obligate intermediate conformation. Here, we provide genetic evidence that open Env conformations can overcome the rhCD4 entry barrier and increase replication of SHIVs in rhesus lymphocytes. Consistent with prior studies, we found that circulating HIV-1 Env do not use rhCD4 efficiently for viral entry. However, using HIV-1 Env with single amino acid substitutions that alter their conformational state, we found that transitions to intermediate and open Env conformation allow usage of physiological levels of rhCD4 for viral entry. We engineered these single amino acid substitutions in the transmitted/founder HIV-1_BG505_ Env encoded by SHIV-BG505 and found that open Env conformation enhances SHIV replication in rhesus lymphocytes. Lastly, CD4-mediated SHIV pull-down, sensitivity to soluble CD4, and fusogenicity assays indicated that open Env conformation promotes efficient rhCD4 binding and viral-host membrane fusion. These findings identify conformational state of HIV-1 Env as a major determinant for rhCD4 usage, viral fusion, and SHIV replication.

**Importance:** Rhesus macaques are critical animal model for preclinical testing of HIV-1 vaccine and prevention approaches. However, HIV-1 does not replicate in rhesus macaques, and thus chimeric simian-human immunodeficiency viruses (SHIVs), which encode HIV-1 envelope glycoproteins, are used as surrogate challenge viruses to infect rhesus macaques for modeling HIV-1 infection. Development of SHIVs encoding envelope from clinically relevant, circulating HIV-1 variants has been extremely challenging as such SHIVs replicate poorly, if at all, in rhesus lymphocytes. This is because most circulating HIV-1 envelope do not use rhesus CD4 efficiently for viral entry. In this study, we identify conformational state of HIV-1 envelope as a key determinant for rhesus CD4 usage, viral-host membrane fusion, and SHIV replication in rhesus lymphocytes.

## Introduction

The ability of viruses to efficiently utilize orthologous entry receptors determines which species can be infected, and this principle is often exploited for developing animal models of viral infection. Rhesus macaques serve as critical animal model for pre-clinical HIV-1 research. However, macaque models are limited by the fact that HIV-1 does not persistently infect macaques. Chimeric simian-human immunodeficiency viruses (SHIVs), constructed by replacing SIV envelope glycoproteins (Env) with that from HIV-1, have been developed as surrogates to study HIV-1 infection in rhesus macaques (1). Development of SHIVs has been a challenging process, mainly because most Env from circulating HIV-1 variants do not use rhesus CD4 (rhCD4) efficiently for viral entry, and, therefore, SHIVs replicate poorly in rhesus lymphocytes and do not establish persistent infection (2). Existing pathogenic SHIVs are a highly selected subset of viruses because they encode Env from lab-adapted and chronic-stage HIV-1 variants, which can mediate entry using rhCD4. Moreover, to increase their pathogenicity SHIVs generally require extensive adaptation to macaques (3–8), at least in part to optimize rhCD4-mediated entry (9). However, adaptation alters the antigenic features of Env (10), limiting the translational utility of adapted SHIVs.

Binding of the metastable HIV-1 Env trimer to CD4 triggers a series of conformational changes in Env i.e., the Env transitions from a functionally ‘closed’ (state 1) to an ‘intermediate’ (state 2) to an ‘open’ (state 3) conformation (11–13). Unliganded Env trimer of primary isolates only infrequently transit from closed to downstream, more open states (11, 14). The conformational dynamics of HIV-1 Env can affect viral tropism by allowing adaptability to cells with different CD4 levels. For example, HIV-1 Env that frequently sample state 2 and state 3 conformations can more efficiently infect cells that express low levels of human CD4 (huCD4) (15, 16). Importantly, the ability of HIV-1 Env to utilize low levels of huCD4 positively correlates with their ability to utilize rhCD4 (17). Based on these insights, we hypothesized that open conformational states of HIV-1 Env can overcome the rhCD4 entry barrier, which can in turn increase replication of SHIVs in rhesus lymphocytes. In this study, we generated HIV-1 Env and SHIVs with single amino acid substitutions that alter the conformational state of Env. Our results show that SHIVs with open Env conformation bind and utilize rhCD4 efficiently and display higher viral fusion and replication in rhesus lymphocytes.

## Results

### Transmitted/founder and early-stage HIV-1 Envs do not use physiological levels of rhCD4 efficiently for viral entry

Env from most circulating HIV-1 isolates from early stages of infection, including the transmitted/founder (T/F) variants, do not use rhCD4 efficiently for viral entry (17). To test our hypothesis, that open Env conformations can overcome the rhCD4 entry barrier and mediate entry using physiological levels of rhCD4 expressed on rhesus lymphocytes, we first generated Cf2Th/syn CCR5 cells that stably express either low or high levels of rhCD4 (rhCD4_LOW_, rhCD4_HIGH_). Importantly, we sorted and selected the rhCD4_LOW_ cells such that their CD4 expression levels is representative of primary rhesus lymphocytes (**Figure 1**). The cells were then infected with GFP-reporter HIV-1 pseudotyped with Env that were obtained from HIV-1 isolates from acute/early stages of infection, including the T/F isolates (**Table 1**). As control, Cf2Th/syn CCR5 cells expressing huCD4 were also infected and facilitated entry by all of these Env (**Figure 2**). In contrast to huCD4-mediated entry, rhCD4_LOW_ cells did not facilitate efficient entry by seven out of eight Env. Even with rhCD4_HIGH_ cells, which express high levels of rhCD4, viral entry was much lower (~2.5–53-fold) than the huCD4. Only subtype D Env 191859 was able to gain entry in rhCD4_LOW_ cells but it was still ~1.7-fold lower than huCD4, highlighting that there are some rare primary Env that can inherently utilize rhCD4. As positive controls, two Env from chronic-stage HIV-1 isolates (JR-FL and BaL) were able to gain entry using both huCD4 and rhCD4 – consistent with the ability of Env from lab-adapted and chronic-stage isolates to utilize rhCD4 for entry (17).

**Figure 1.**
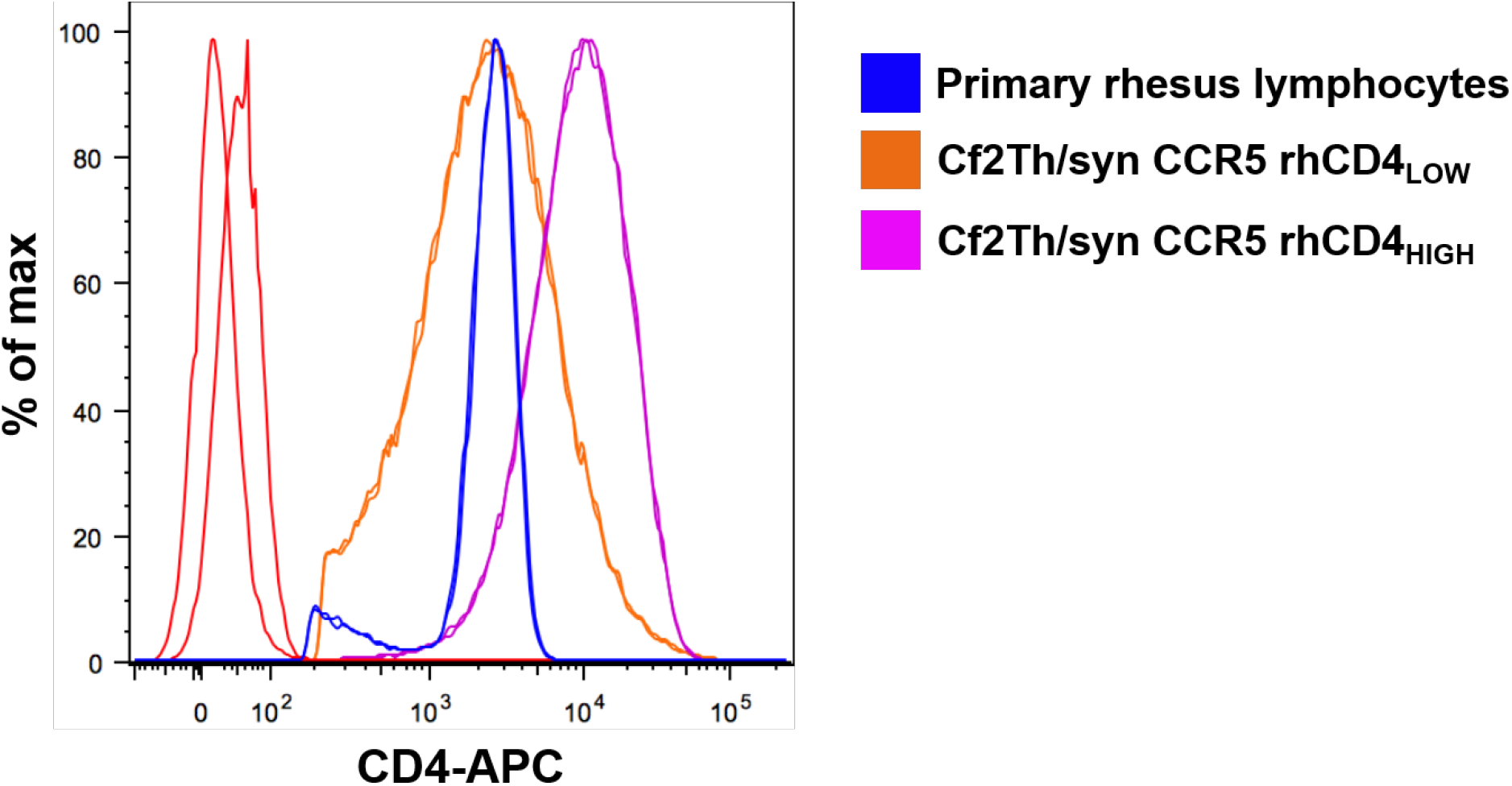
Expression levels of rhCD4 stably introduced into Cf2Th/syn CCR5 cells. Expression levels of CD4s on primary rhesus macaque lymphocytes, and rhCD4_LOW_ and rhCD4_HIGH_ Cf2Th/syn CCR5 cell lines as measured by flow cytometry using an APC-conjugated anti-CD4 antibody. The histograms represent the expression of CD4 on the cell surface. The data are representative of two independent experiments.

**Figure 2.**
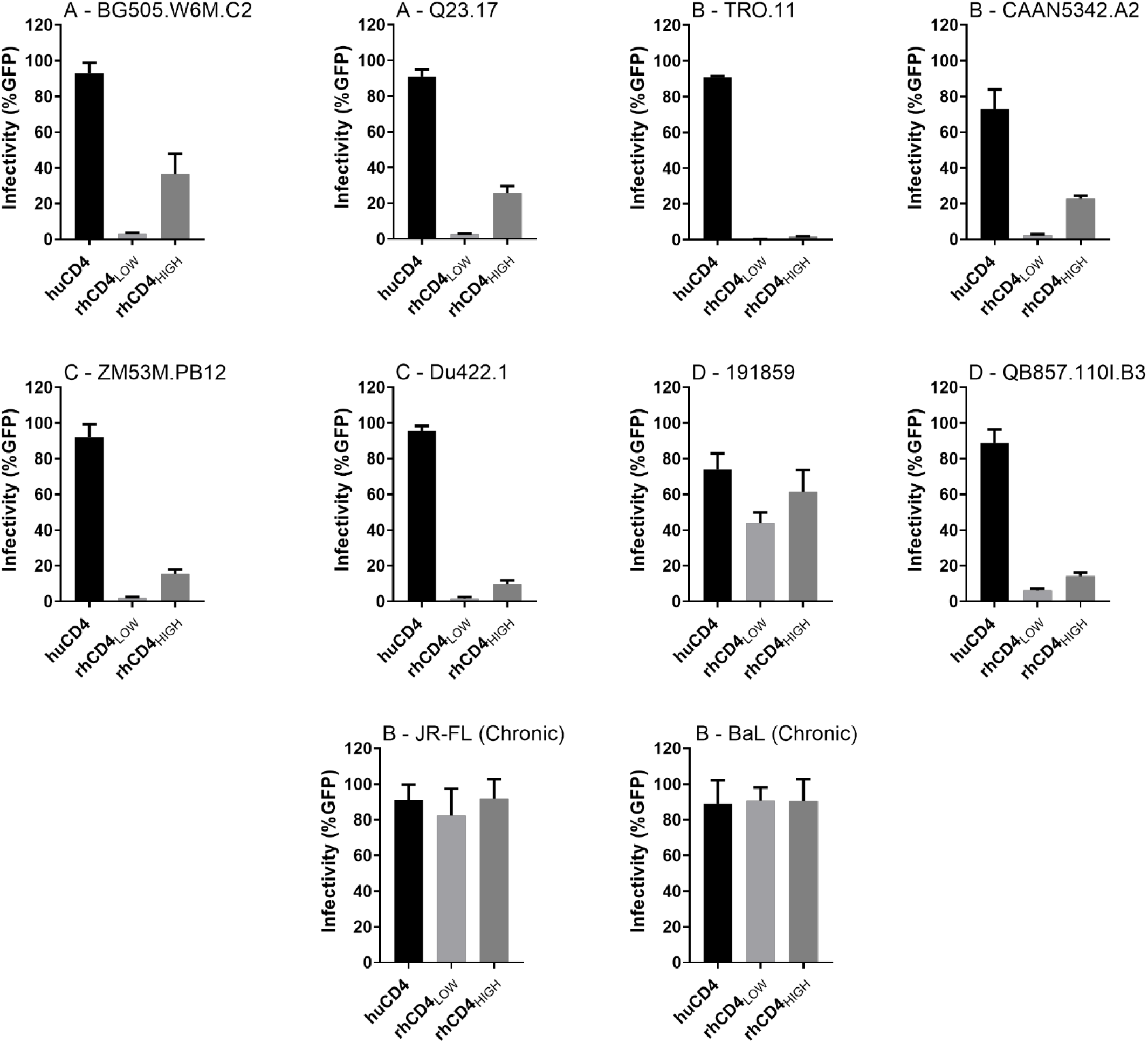
Ability of transmitted/founder and early-stage HIV-1 Env to use physiological levels of rhCD4 for viral entry. Cf2Th/syn CCR5 cells expressing human or rhesus CD4 (indicated along the x-axis) were infected with HIV-1_Q23ΔEnvGFP_ pseudotyped with indicated Env clones. Infection was measured by flow cytometry as percentage of GFP-positive cells 48 h post-infection. Graphs indicate percentage of infected cells for indicated Env at multiplicity of infection of 1. Env clones are labeled at the top, with first letter indicating the subtype. Bars represent the average of three independent experiments. Error bars represent standard deviations.

**Table 1:** Transmitted/founder and early-stage Env clones used in this study.

| Envelope clone | Subtype | Mode of transmission<sup>a</sup> | Source<sup>b</sup> | GenBank accession no. | Time post infection (weeks) |
| --- | --- | --- | --- | --- | --- |
| BG505.W6M.C2 | A | M-C | PBMC | DQ208458 | 6 |
| Q23.17 | A | M-F | PBMC | AF004885 | 11 |
| TRO.11 | B | M-M | ccPBMC | AY835445 | 4 |
| CAAN5342.A2 | B | M-M | Plasma | AY835452 | <12 |
| Du422.1 | C | M-F | ccPBMC | DQ411854 | 8 |
| ZM53M.PB12 | C | F-M | PBMC | AY423984 | <12 |
| QB857.110I.B3 | D | M-F | PBMC | FJ866138 | 16 |
| 191859 | D | M-F | Plasma | JX203064 | Fiebig I |

<sup>a</sup> M-C: Mother to Child; M-F: Male to Female; M-M: Male to Male; F-M: Female to Male.
<sup>b</sup> PBMC: Env cloned from uncultured PBMCs isolated directly from patient; ccPBMC: Patient PBMCs (or virus from these PBMCs) underwent short-term coculture with PBMCs from HIV-1-negative donors to amplify virus before cloning; Plasma: Env cloned from virion RNA in plasma isolated directly from patient.

### Transitions to intermediate and open Env conformations allow usage of rhCD4 for viral entry

Recent studies have suggested that Env of most primary HIV-1 isolates exist in a closed conformation and only infrequently transit to downstream, more open conformations (11, 13, 18, 19). The ability of unliganded or CD4-bound Env to transit to downstream states can affect viral tropism by allowing adaptability to cells with different CD4 levels (15, 16). Thus, we sought to determine how the conformational states of HIV-1 Env affects its ability to utilize rhCD4 for viral entry. We took advantage of the fact that single amino acid substitutions in the gp120 domain of Env can alter its conformational state (15, 16). For example, L193A substitution in the V1/V2 loop of gp120 allows Env to populate the state 2 conformation. I423A substitution in the β20-β21 element of gp120 allows Env to populate the state 3 conformation. We engineered L193A and I423A substitutions in three different Env representing a T/F (BG505.W6M.C2), an acute stage (ZM53M.PB12), and a chronic stage (JR-FL) isolate. For the two early-stage Env, I423A substitution mediated entry in rhCD4_LOW_, rhCD4_HIGH_, and huCD4 cells with similar efficiency, whereas the L193A substitution displayed lower efficiency of entry in rhCD4_LOW_ cells when compared to huCD4 and rhCD4_HIGH_ (**Figure 3A and 3B**). As control, the related wild type (WT) Env utilized huCD4 efficiently but did not gain efficient entry even in cells that express high levels of rhCD4. The chronic-stage JR-FL Env was able to gain entry using both huCD4 and rhCD4 – independent of Env conformational state or rhCD4 expression levels (**Figure 3C**). Similar patterns of rhCD4 usage were observed at different multiplicities of infection indicating that the ability of open Env conformations to use physiological levels of rhCD4 is independent of virus input (**Figure S1**). Thus, our results suggest that single amino acid substitutions in Env that promote transitions to intermediate and open conformations allow usage of physiological levels of rhCD4 for viral entry.

**Figure 3.**
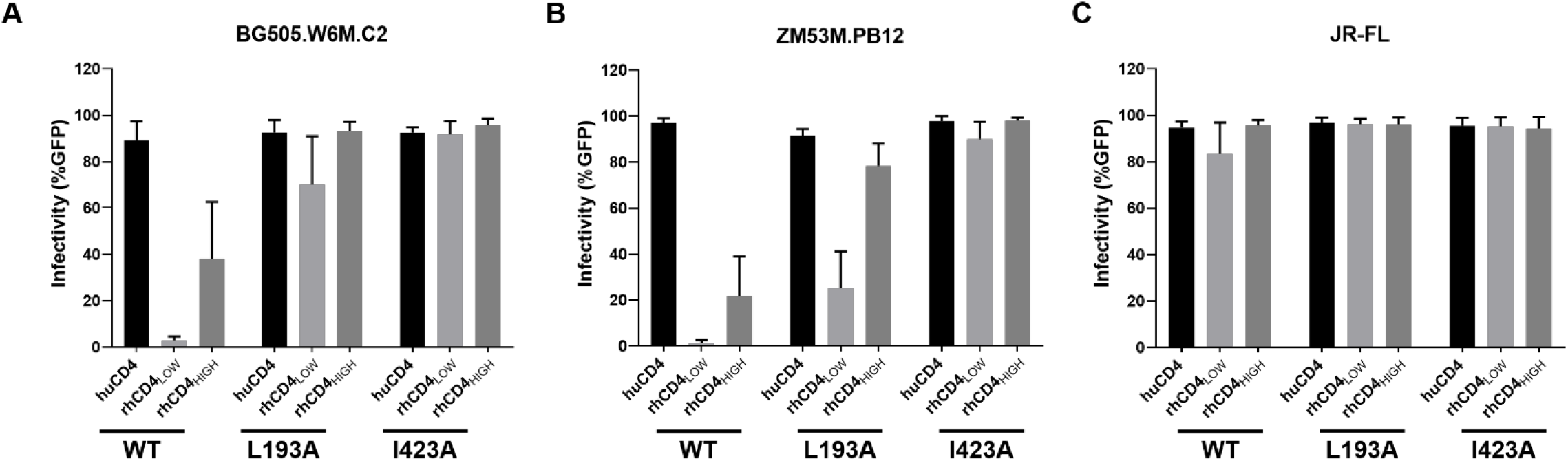
Effect of Env conformation on usage of rhCD4 for viral entry. Cf2Th/syn CCR5 cells expressing human or rhesus CD4 (indicated along the x-axis) were infected with HIV-1_Q23ΔEnvGFP_ pseudotyped with **(A)** BG505.W6M.C2, **(B)** ZM53M.PB12, and **(C)** JR-FL Env clone. Infection was measured by flow cytometry as percentage of GFP-positive cells 48 h post-infection. Graphs indicate percentage of infected cells for wild-type (‘WT’), L193A, and I423A Env variants at multiplicity of infection of 1. Bars represent the average of three independent experiments. Error bars represent standard deviations.

### Open Env conformations promote replication of SHIV-BG505 in rhesus lymphocytes

Next, we sought to determine whether improved rhCD4 usage by Env that are in intermediate and open conformational states translate into enhanced replication of T/F SHIV in rhesus lymphocytes. For this purpose, we introduced the L193A or I423A substitution in the HIV-1_BG505_ T/F Env encoded by SHIV-BG505. Importantly, we also included SHIV-BG505 with A204E substitution in the C1 region of gp120 as a benchmark control for replication. The A204E adaptive mutation, previously identified through serial passage of primary HIV-1 Env in immortalized macaque CD4^+^ T-lymphocytes (20), was sufficient to mediate viral entry in rhCD4_LOW_ cells at levels comparable to huCD4 (**Figure S2**). More importantly, the A204E substitution induces conformational changes in the Env that opens up the Env trimers on the surface of virions (10), which likely explains its ability to utilize physiological levels of rhCD4. We measured the ability of these SHIVs to replicate in the immortalized rhesus macaque CD4^+^ T-lymphocytes (21). As expected, the replication of SHIV encoding wild-type Env (WT SHIV), which does not utilize rhCD4 efficiently, declined over the 15-day time course indicating that it does not replicate in rhesus cells (**Figure 4A and 4B**). SHIVs encoding I423A or A204E substitutions replicated to significantly higher levels over the 15-day time course when compared to WT SHIV. Interestingly, SHIV L193A, which encodes Env with an intermediate open conformation that allows it to use rhCD4 better than the WT Env (**Figure 3A**), did not display significantly improved growth kinetics when compared to WT SHIV. Based on these findings, we conclude that open Env conformations, such as those conferred by I423A and A204E mutations, promote replication of SHIV-BG505 in rhesus lymphocytes.

**Figure 4.**
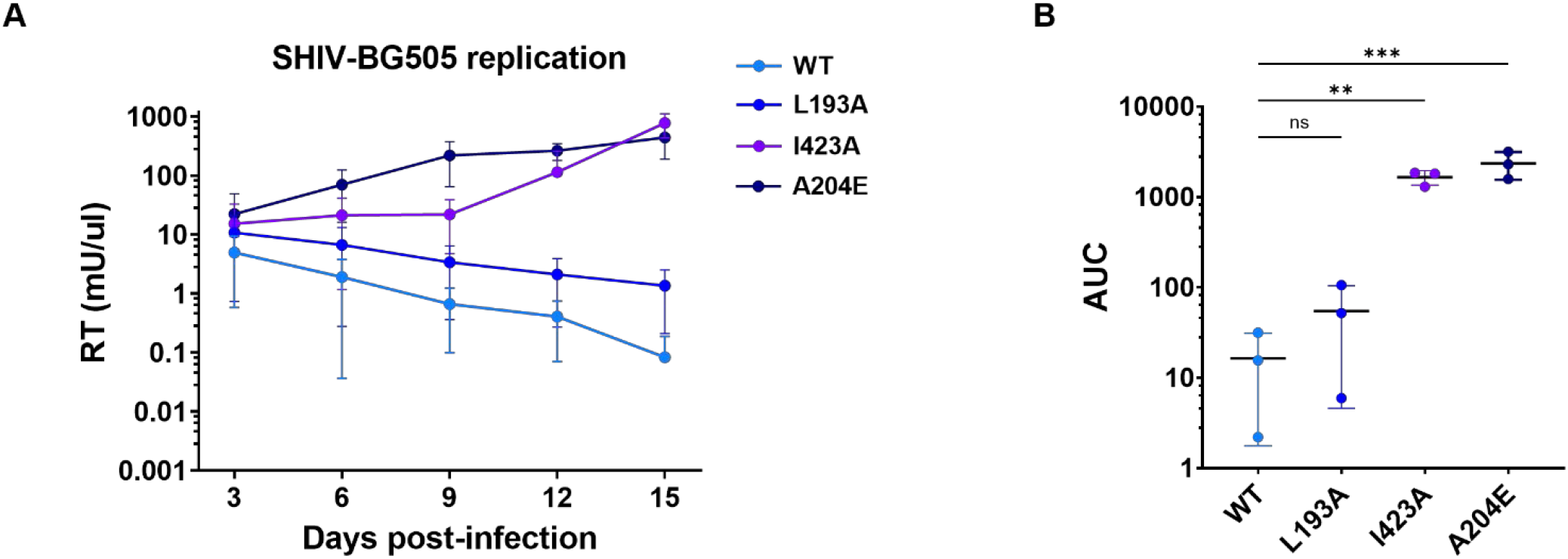
Effect of Env conformation on SHIV replication. **(A)** Replication kinetics of SHIV-BG505 variants in rhesus macaque 221 T-lymphocytes over a 15-day time course. Reverse transcriptase (RT) activity in viral supernatants is plotted vs. days post-infection. The key at the right of the graph indicates the identity of Env variant (wild-type ‘WT’ or indicated amino acid mutation) encoded by each SHIV. Each data point represents the average of three independent experiments, performed in duplicate. Error bars represent standard deviations. **(B)** Area under the curve (AUC) for indicated SHIV variants (x-axis) determined from the replication curves shown in (A). AUC values were compared to SHIV-BG505 WT using one-way analysis of variance (ANOVA) followed by Dunnett’s multiple comparisons test. *** *p* = 0.0004; ** *p* = 0.004; ns, not significant.

### Open Env conformations promote CD4 binding and viral fusion

Finally, we examined whether open Env conformations promote SHIV replication by enhancing rhCD4 binding and viral fusion. We evaluated the effects of Env conformation on rhCD4 binding by two independent approaches. First, we performed neutralization assays to measure the ability of soluble rhCD4 to bind functional Env trimers on SHIVs and compete with receptor binding to inhibit viral entry. WT SHIV was least sensitive to rhCD4 inhibition indicating that it does not efficiently bind rhCD4 and interfere with receptor binding (**Figure 5A**). SHIV L193A and SHIV I423A, which are stabilized in State 2 and State 3 Env conformation, respectively, were ~3-fold and ~30-fold more sensitive to rhCD4 inhibition when compared to WT SHIV, suggesting that progressive opening of the Env increases rhCD4 binding. SHIV A204E was most sensitive (~1500-fold) to rhCD4 inhibition suggesting that it has the highest binding for rhCD4, likely attributable to ‘more open’ conformation of its Env trimers. Similar neutralization trends, but with ~2-10-fold more potent inhibition, were observed with soluble huCD4 indicating that HIV-1 Env encoded by these SHIVs bind huCD4 with higher affinity than rhCD4 (**Figure S3A**). Second, we employed a qualitative affinity pull-down assay to determine the binding of rhCD4 to Env expressed on infectious SHIVs. Virions bound to His-tagged rhCD4 were affinity precipitated and their Env levels were measured by immunoblotting. Consistent with the results of the neutralization assay, progressive opening of the Env resulted in more virions binding to rhCD4; with highest binding observed in case of SHIV A204E (**Figure 5B**). Similar binding patterns were also observed when pull-downs were performed with huCD4 (**Figure S3B**). Lastly, we investigated how Env conformation affects fusion of SHIVs with rhesus lymphocytes using a SHIV fusion assay. WT SHIVs, which did not efficiently bind rhCD4, were not fusogenic with the target cells (**Figure 5C and 5D**). SHIV L193A and SHIV I423A, which are stabilized in State 2 and State 3 Env conformation, respectively, were ~7-fold and ~25-fold more fusogenic when compared to WT SHIV, suggesting that fusogenicity increases with progressive opening of the Env. SHIV A204E was the most fusogenic with ~87-fold higher fusion than WT SHIV. Taken together, our results demonstrate that open Env conformations, such as those conferred by I423A and A204E mutations, increase binding of Env to rhCD4 and subsequent SHIV-host membrane fusion.

**Figure 5.**
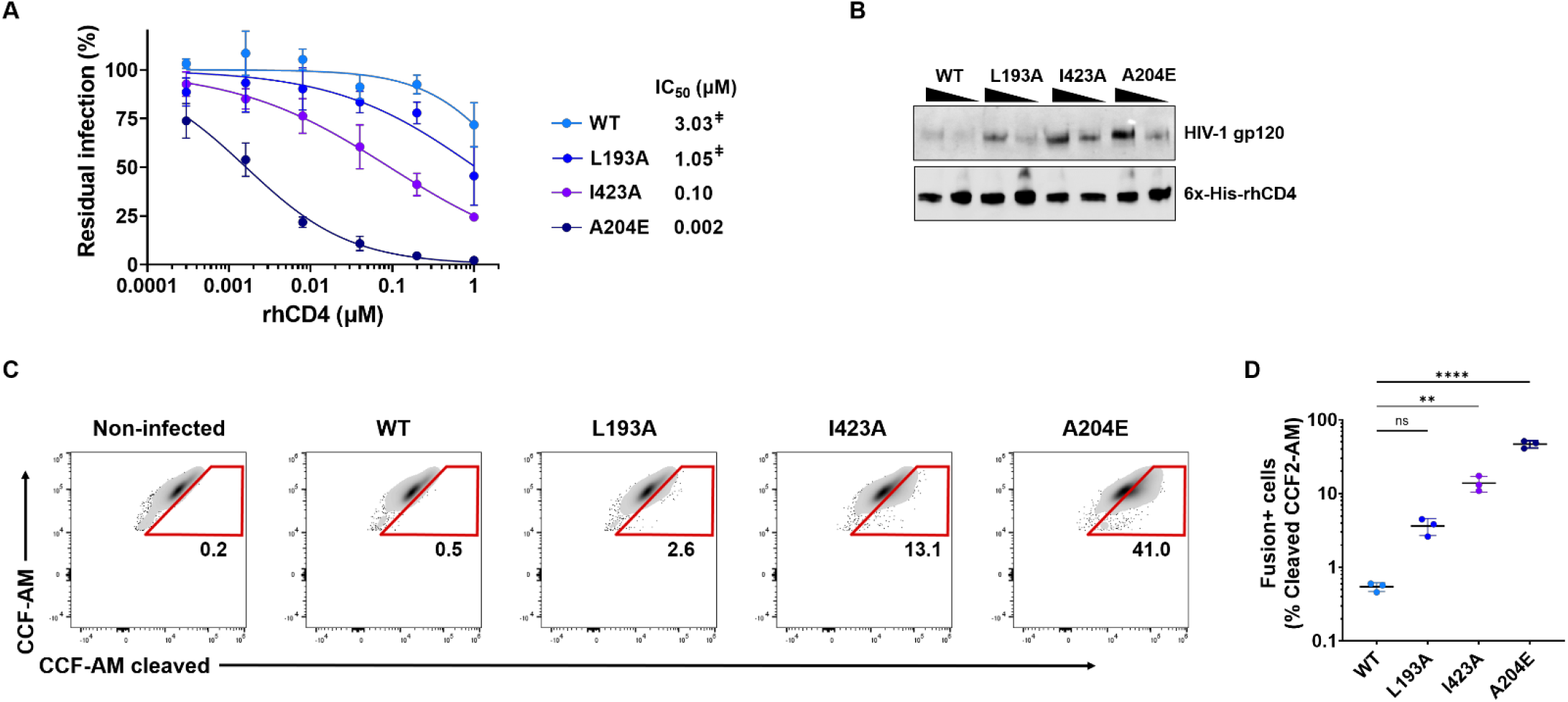
Effect of Env conformation on rhCD4 binding and viral fusion. **(A)** Sensitivity of SHIV-BG505 to neutralization by soluble rhCD4. Neutralization curves of the indicated SHIV variants were generated by plotting percent residual infection (y-axis) against rhCD4 concentration (μM, x-axis). Each data point represents the average of two independent experiments, performed in duplicate. Error bars represent standard deviations. The calculated IC_50_s are shown. Where IC_50_ values were above the highest tested concentration, the extrapolated concentration is indicated by ^‡^. **(B)** Binding of SHIV-BG505 to rhCD4. Western blot analysis for affinity pull-down of His-tagged rhCD4 with increasing amounts (400 and 800 mU of RT) of indicated SHIV virions. Immunoblotting performed using anti-HIV-1 gp120 and anti-6x-His antibodies. **(C)** Representative flow cytometry plots indicating percent viral fusion. Fusion of indicated SHIV-BG505 with rhesus macaque 221 T-lymphocytes was measured as percentage of cells with cleaved CCF-AM substrate. The identity of Env variant (wild-type ‘WT’ or indicated amino acid mutation) encoded by each SHIV is indicated above the plots. **(D)** Graph indicate percent viral fusion for indicated SHIV variants (x-axis). Data represents the average of three independent experiments with individual data points shown as circles. Error bars represent standard deviations. Percent viral fusion were compared to SHIV-BG505 WT using one-way analysis of variance (ANOVA) followed by Dunnett’s multiple comparisons test. **** *p* < 0.0001; ** *p* = 0.0024; ns, not significant.

## Discussion

Here, we provide first genetic evidence that conformational state of HIV-1 Env governs rhCD4 usage. Using HIV-1 Env with single amino acid substitutions that alter their conformational state, we demonstrated that transitions to intermediate and open Env conformations allow usage of physiological levels of rhCD4 for viral entry, which is otherwise a suboptimal receptor for entry. Moreover, by introducing these changes in isogenic SHIV backbone, we found that open Env conformations promote rhCD4 binding, fusion-mediated cell entry, and replication in rhesus lymphocytes.

Thermodynamically, most primary HIV-1 Env are in a high free energy, closed conformation. Single-molecule fluorescence resonance energy transfer studies have demonstrated that either spontaneously or upon CD4 binding, Env transitions from closed (state 1) to open conformation (state 3) through a functional intermediate (state 2) (11). Multiple amino acid residues restrain Env in state 1 and CD4 binding triggers Env transitions from state 1 to downstream, lower-energy states (15, 16, 22–25). For example, L193A and I423A substitution releases the state 1 restraints and stabilizes Env in State 2 and State 3 conformation, respectively. Our findings of improved rhCD4 usage by Envs bearing L193A and I423A substitutions suggest that transitions to intermediate and open conformations advance HIV-1 Env on the entry pathway and likely facilitate Env engagement with a suboptimal rhCD4 receptor. Although Env with L193A substitutions, which are in intermediate open conformation, utilized rhCD4 better than WT Env, SHIV encoding L193A substitution did not display significantly improved viral fusion and replication. In contrast, SHIVs encoding Env with I423A and A204E substitutions, which are in open conformation, displayed significantly increased rhCD4 engagement, viral fusion, and replication. These results indicate that open Env conformations, such as those conferred by I423A and A204E mutations, promote rhCD4 binding, viral fusion, and replication of SHIV in rhesus lymphocytes. Collectively, these findings suggest that modulating the conformational dynamics of viral Env by altering select amino acid residues can help overcome the cross-species entry barrier, which in turn facilitates replication and adaptation of the virus in a new host species.

While our findings suggest that downstream, open Env conformations allow usage of rhCD4, it also highlights that most T/F and early-stage HIV-1 Envs likely have an inherently low propensity to sample downstream, open conformations. For instance, only one of the eight early-stage Env tested was able to use physiological levels of rhCD4 for viral entry. This explains why development of SHIVs encoding T/F and early-stage Envs has been challenging and often require extensive adaptation in human and/or rhesus lymphocytes – process that selects for open Env conformation (2, 10, 20). Consistent with this notion, SHIV-BG505 encoding A204E adaptive mutation, previously identified through *in vitro* evolution experiments (20), displayed highest replication in rhesus lymphocytes; viral-host membrane fusion; and sensitivity to soluble CD4. These findings also offer an explanation as to why some existing pathogenic SHIVs derived using primary HIV-1 Envs display high sensitivity to soluble CD4 (26–28) – a proxy measure for open Env conformation where the CD4 binding site is exposed.

In summary, our findings have helped identify a key parameter for future design of SHIVs – conformational state of Env. Considerable efforts are being made to engineer SHIVs that encode circulating HIV-1 Env; utilize rhCD4 for entry; replicate in rhesus macaques without extensive adaptation; and retain as much of the biological characteristics of HIV-1 Env as possible (29–31). Thus, it will be of interest to identify and define the minimal open conformational state of HIV-1 Env that affords usage of rhCD4 and replication of SHIVs in rhesus macaques with minimal impact on Env trimer structure and antigenic profile. Based on the findings of this study, the following considerations could be useful for SHIV design: 1) selection of circulating HIV-1 Env with high inherent propensity to sample downstream, open conformations; and 2) engineering changes in Env that lower free energy needed for transition to open conformations without significantly altering its antigenicity.

## Materials and Methods

### Cells, envelope clones, plasmids, proteins

HEK293T (ATCC CRL-3216), HeLa TZM-bl (32) (NIH AIDS Reagent program catalog no. 8129), and Cf2Th/syn CCR5 (33) (NIH AIDS Reagent program catalog no. 4662) cells were cultured in Dulbecco’s modified eagle medium (DMEM, Gibco) supplemented with 10% fetal bovine serum (FBS, Sigma), 2 mM L-glutamine (Gibco), and 1x Penicillin-Streptomycin (Gibco) (complete DMEM). Cf2Th/syn CCR5, which are engineered to express human CCR5, were further supplemented with 400 μg/ml of Geneticin (Gibco) to maintain CCR5 expression. Immortalized rhesus macaque 221 T-lymphocytes (21) were cultured in Iscove’s modified Dulbecco’s medium (IMDM) supplemented with 10% FBS, 2 mM L-glutamine, and 1x Penicillin-Streptomycin, and 100 U/ml of interleukin-2 (Roche) (complete IMDM). Rhesus macaque peripheral blood mononuclear cells (PBMCs) were isolated from whole blood (Washington National Primate Research Center) from two independent donors using human erythrocyte lysing kit (R&D Systems) and 95% Lymphoprep (Stemcell Technologies) following manufacturer’s protocols.

The following Env clones from early HIV-1 infections were used: BG505.W6M.C2; Q23.17; TRO.11; CAAN5342.A2; Du422.1; ZM53M.PB12; QB857.110I.B3; and 191859. As control, two Env clones (JR-FL and BaL.01) of chronic HIV-1 strains were used. The following Env clones encoding the L193A, I423A or A204E mutations were used: BG505.W6M.C2; ZM53M.PB12; and JR-FL.

β-lactamase (BlaM) gene fused to the N-terminus of SIVmac239 Vpr (Genbank: M33262) separated by six-glycine-one-lysine linker was synthesized as a DNA fragment (Integrated DNA Technologies), digested, and ligated into pcDNA3.1 (Invitrogen) using KpnI and NotI restriction sites to generate pcDNA3.1-Blam-SIVmac239 Vpr plasmid. All plasmids generated in this study were verified by Sanger DNA sequencing.

Soluble rhCD4 (Met1-Trp390; catalog no. 90274-C08H) and huCD4 (Met 1-Trp 390; catalog no. 10400-H08H) proteins with C-terminus 6x-His-tags were purchased from Sino Biological.

### Pseudovirus and SHIV production

Green fluorescent protein (GFP) reporter pseudoviruses were generated as described previously (34). Briefly, HEK293T cells were cotransfected with 4 μg of Env-deficient HIV-1 proviral plasmid (Q23ΔEnvGFP) and 2 μg of HIV-1 Env clone of interest using Fugene 6 transfection reagent (Roche) following manufacturer’s protocol. Replication-competent SHIV stocks were generated as described previously (35). The viral titer of each pseudovirus and SHIV stock was determined by infecting TZM-bl cells and staining for β-galactosidase activity 48 hours post-infection (32).

### Generation of stable cell lines

Cf2Th/syn CCR5 cells stably expressing rhCD4 were generated using methods described previously (36). Briefly, retroviral pseudoviruses were generated in HEK293T cells by cotransfecting pLPCX-rhCD4 (retroviral vector encoding rhCD4), pJK3 (MLV-based packaging plasmid), and pMD.G (vesicular stomatitis virus glycoprotein [VSV-G] plasmid) at a ratio of 1:1:0.5 using Fugene 6 transfection reagent following manufacturer’s protocol. Forty-eight hours post-transfection, the pseudoviruses were concentrated and used to transduce 10^5^ Cf2Th/syn CCR5 cells. Transduced cells were cultured in complete DMEM supplemented with 400 μg/ml of Geneticin (to maintain CCR5 expression) and 2 μg/ml of puromycin (to select for CD4 expression, Sigma). The drug selected cells with low (rhCD4_LOW_) or high (rhCD4_HIGH_) levels of rhCD4 expression were obtained by sorting the cells on a FACSAria II cell sorter (BD Biosciences) using an allophycocyanin (APC)-conjugated CD4 monoclonal antibody (BD Biosciences catalog no. 551980) using previously described CD4 staining method (17). The rhCD4_LOW_ cells were sorted by gating on the CD4-expressing cell population that overlapped with the CD4-stained rhesus macaque PBMCs. The rhCD4_HIGH_ cells were sorted by gating on the top 20% of CD4-expressing cell population. Cf2Th/syn CCR5 cells stably expressing huCD4 have been described previously (36).

### CD4 infectivity assay

Infection of Cf2Th/syn CCR5 cells expressing huCD4, rhCD4_LOW_, and rhCD4_HIGH_ with GFP reporter pseudoviruses was performed as described previously (36). Briefly, cells were infected at a multiplicity of infection (MOI) of 0.1, 0.25, and 1 in the presence of 10 μg/ml of DEAE-dextran by spinoculation at 1200 × g for 90 minutes. After 48 hours, cells were harvested, fixed in 2% paraformaldehyde, washed twice, and resuspended in fluorescence-activated cell sorter (FACS) buffer (1x phosphate-buffered saline (PBS), 1% FBS, 1 mM EDTA). Cells were analyzed for GFP expression on Attune NxT flow cytometer (Life Technologies). The data from ~10^4^ cells were analyzed using FlowJo version 10.7.1.

### Construction of SHIV proviral clones

Full-length proviral SHIV plasmid encoding BG505.W6M.B1 Env (37) with A204E mutation (SHIV-BG505 A204E) has been described previously (38). EcoRV-MfeI region of BG505.W6M.B1 Env (wild type, WT) and its L193A and I423A variants were synthesized as DNA fragments (Twist Biosciences). SHIV-BG505 WT, SHIV-BG505 L193A, and SHIV-BGB505 I423A proviral clones were generated by digesting and ligating the DNA fragments into the SHIV-BG505 A204E proviral plasmid using EcoRV and MfeI restriction sites. The generated SHIV proviral plasmids were verified by DNA sequencing.

### SHIV replication time course

Replication of SHIVs was assessed as described previously (35). Briefly, 4×10^6^ 221 T-lymphocytes were infected at a MOI of 0.02 by spinoculation at 1200 × g for 90 minutes at room temperature. After spinoculation, cells were washed four times with 1 ml of complete IMDM, re-suspended in 5 ml of complete IMDM and plated in one well of a 6-well plate. Every three days, two-third of the cultures were harvested and replenished with fresh, complete IMDM. Viral supernatants were collected from the harvested cultures by pelleting at 650 × g for 5 minutes at room temperature. RT activity in viral supernatants was measured using the RT activity assay.

### SHIV neutralization assay

Neutralization of SHIVs with rhCD4 and huCD4 were performed as described previously (16) but using TZM-bl target cells. The calculated half-maximal inhibitory concentration (IC_50_) values represent the soluble CD4 concentration in μM at which 50% of the virus was neutralized.

### CD4–SHIV binding assay

Affinity pull-down assays were performed using Ni Sepharose 6 Fast Flow resin (Cytiva Lifesciences); His-tagged rhCD4 or huCD4; and SHIV stocks to assess binding of CD4 to SHIVs. 20 μl of Ni-resin was equilibrated in binding buffer (50 mM HEPES pH 7.5, 250 mM NaCl, 50 mM imidazole, and 2 mM β-mercaptoethanol) by washing three times (10,000 × g for 1 minute) with 100 μl of binding buffer. Equilibrated Ni-resin was incubated for 10 minutes with 1 μg of His-CD4 in 20 μl of binding buffer. Binding reactions were setup by adding equal amounts of indicated SHIV virions (equivalent to 800 and 400 mU of RT) in 160 μl of binding buffer and incubated for 1 hour at room temperature, under rotation. Reactions were then washed three times (10,000 × g for 1 minute) with 100 μl of binding buffer to remove unbound CD4/virions. The resulting CD4–virion complexes bound to the Ni-resin were extracted using 4x Laemmli sample buffer (BioRad), heated at 95°C for 5 minutes, and subjected to SDS-PAGE/Western blotting analysis. Standard Western blotting procedures were used with the following antibodies: HIV-1 gp120 (NIH AIDS Reagent program catalog no. 288) and 6x-His Tag (ThermoFisher catalog no. MA1-21315-HRP).

### SHIV fusion assay

The HIV-1 β-lactamase-Vpr (BlaM-Vpr) virus fusion assay described previously (39, 40) was modified to evaluate SHIV fusion. Briefly, BlaM-Vpr was modified by swapping the HIV-1 Vpr with SIVmac239 Vpr to retain the cognate Gag p6-SIVmac Vpr interaction necessary for efficient virion incorporation of Vpr (41). SHIVs containing Blam-SIVmac239 Vpr fusion protein were generated by co-transfecting HEK293T cells with 4.5 μg of proviral SHIV plasmid and 1.5 μg of pcDNA3.1-Blam-SIVmac239 Vpr plasmid using Fugene 6 transfection reagent following manufacturer’s protocol. Forty-eight hours post-transfection, virus-containing supernatant was harvested, passed through a 0.2 μm sterile filter, concentrated ~10-fold using Amicon Ultracel 100 kDa filters (Millipore), aliquoted and stored at −80°C. RT activity of viral stocks was measured using the RT activity assay. 10^5^ 221 T-lymphocytes in 100 μl of complete IMDM were infected with Blam-SIVmac239 Vpr-containing SHIVs equivalent to 720 mU of RT by spinoculation at 1200 × g for 90 minutes followed by incubation at 37°C and 5% CO_2_ for one hour. Fusion-mediated SHIV entry was quantified by monitoring the conversion of fluorescent BlaM CCF2-AM substrate dye as described. After infection, cells were washed once with 300 μl of cold CO_2_-independent media (Gibco) without FBS and resuspended in 100 μl of CO_2_-independent media supplemented with 10% FBS. Cells were incubated with CCF2-AM substrate (LiveBLAzer CCF2-AM Kit, Invitrogen) following manufacturer’s protocol in the presence of 1.8 mM Probenecid (Sigma) for 2 hours. Cells were washed three times with 300 μl of cold CO_2_-independent media without FBS, once with 1x PBS, fixed with 200 μl of 2% paraformaldehyde, washed once with 1x PBS, and resuspended in 400 μl of FACS buffer. The fluorescence of cleaved CCF2-AM substrate was measured on Attune NxT flow cytometer and data were analyzed using FlowJo version 10.7.1.

### Reverse transcriptase activity assay

RT activity assay was performed as described previously (40, 42) with minor modifications. Briefly, 5 μl of viral supernatant, viral stock or RT standard was lysed in 5 μl of 2x lysis buffer (100 mM Tris HCl pH 7.4, 50 mM KCl, 0.25% Triton X-100, 40% glycerol) in the presence of 4U RNaseOUT (Invitrogen) for 10 minutes at room temperature. Viral lysate was diluted 1:10 by adding 90 μl of nuclease-free water (Life Technologies). qRT-PCR reactions were prepared by mixing 9.6 μl of diluted viral lysate with 10.4 μl of reaction mix containing 10 μl of 2x Maxima SYBR Green/ROX qPCR Master Mix (ThermoFisher), 0.1 μl of 4U/μl RNaseOUT, 0.1 μl of 0.8 μg/μl MS2 RNA template (Roche), and 0.1 μl each of 100 μM forward 5’-TCCTGCTCAACTTCCTGTCGAG-3’ and reverse 5’-CACAGGTCAAACCTCCTAGGAATG-3’ primers. qRT-PCR was performed using a QuantStudio 3 Real-Time PCR machine (Applied Biosystems). Viral titers were calculated from a standard curve generated using recombinant reverse transcriptase (Millipore catalog no. 382129).

---

**Figure S1.**
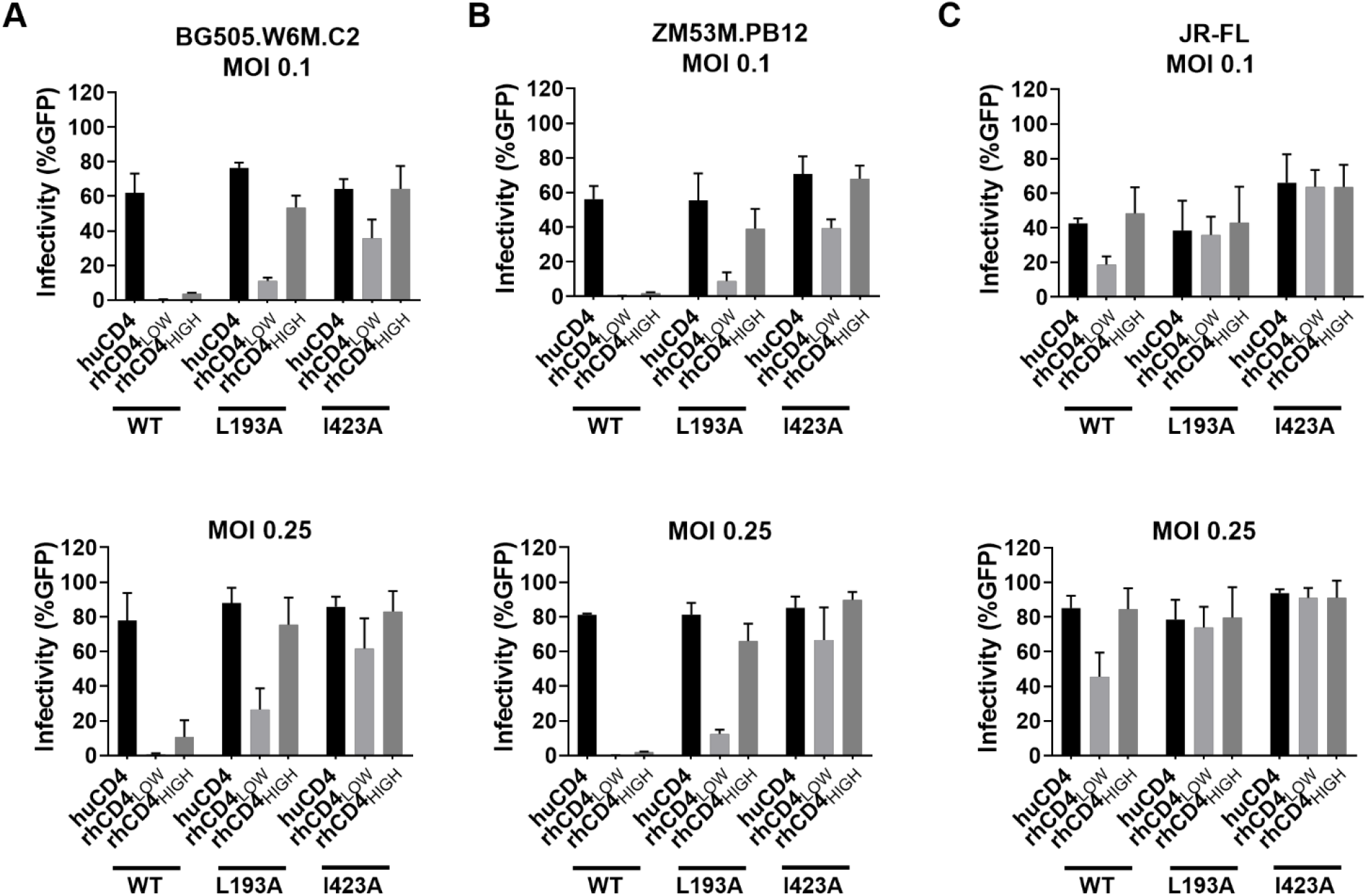
Effect of virus input on rhCD4 usage by HIV-1 Envs in different conformations. Cf2Th/syn CCR5 cells expressing human or rhesus CD4 (indicated along the x-axis) were infected with HIV-1_Q23ΔEnvGFP_ pseudotyped with **(A)** BG505.W6M.C2, **(B)** ZM53M.PB12, or **(C)** JR-FL Env clone. Infection was measured by flow cytometry as percentage of GFP-positive cells 48 h post-infection. Graphs indicate percentage of infected cells for wild-type (‘WT’), L193A, and I423A Env variants at different multiplicity of infection (‘MOI’, *top panel* MOI 0.1, *bottom panel* MOI 0.25). Bars represent the average of three independent experiments. Error bars represent standard deviations.

**Figure S2.**
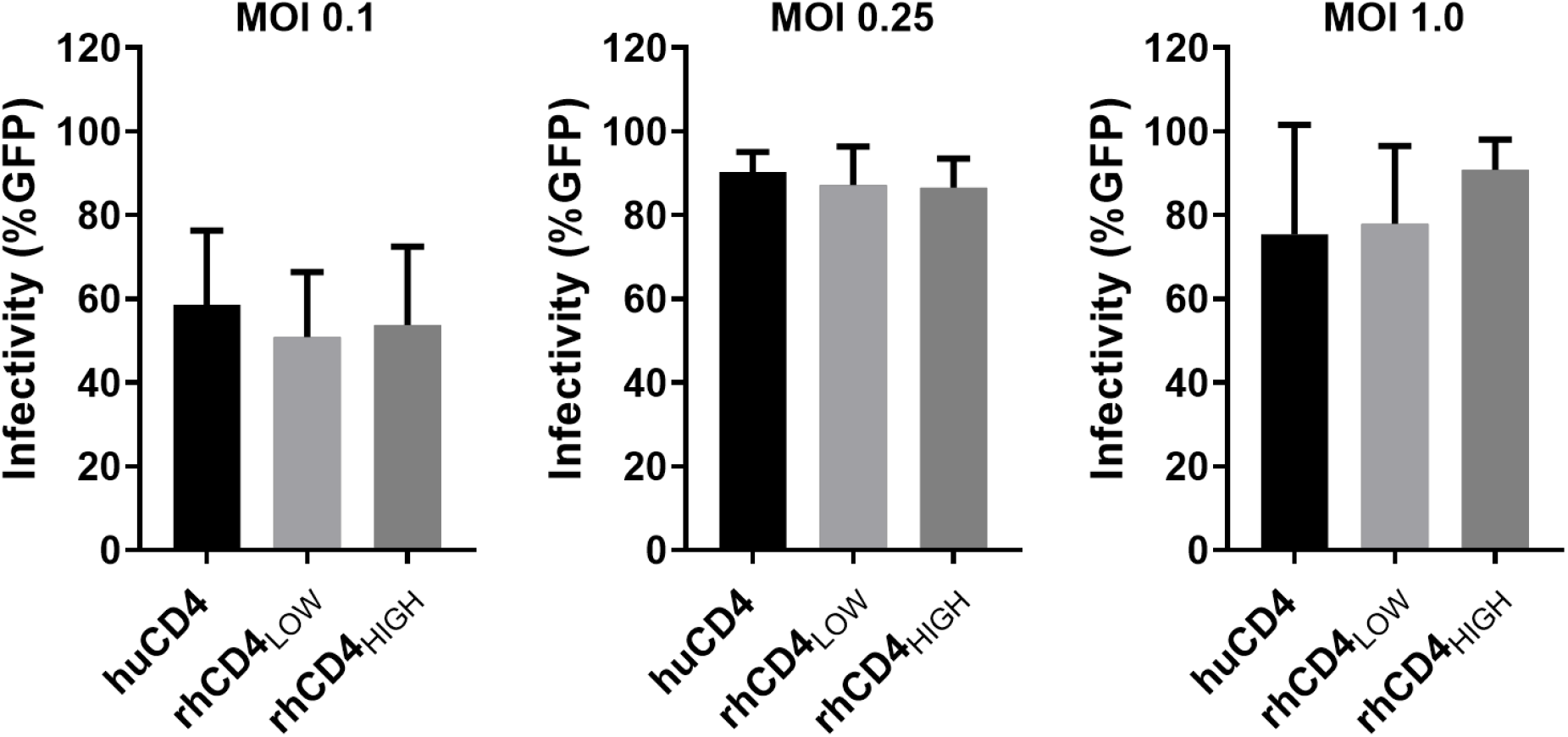
Ability of HIV-1 Env with A204E substitution to use physiological levels of rhCD4 for viral entry. Cf2Th/syn CCR5 cells expressing human or rhesus CD4 (indicated along the x-axis) were infected with HIV-1_Q23ΔEnvGFP_ pseudotyped with BG505 Env encoding A204E mutation. Infection was measured by flow cytometry as percentage of GFP-positive cells 48 h post-infection. Graphs indicate percentage of infected cells at indicated multiplicity of infection (‘MOI’). Bars represent the average of three independent experiments. Error bars represent standard deviations.

**Figure S3.**
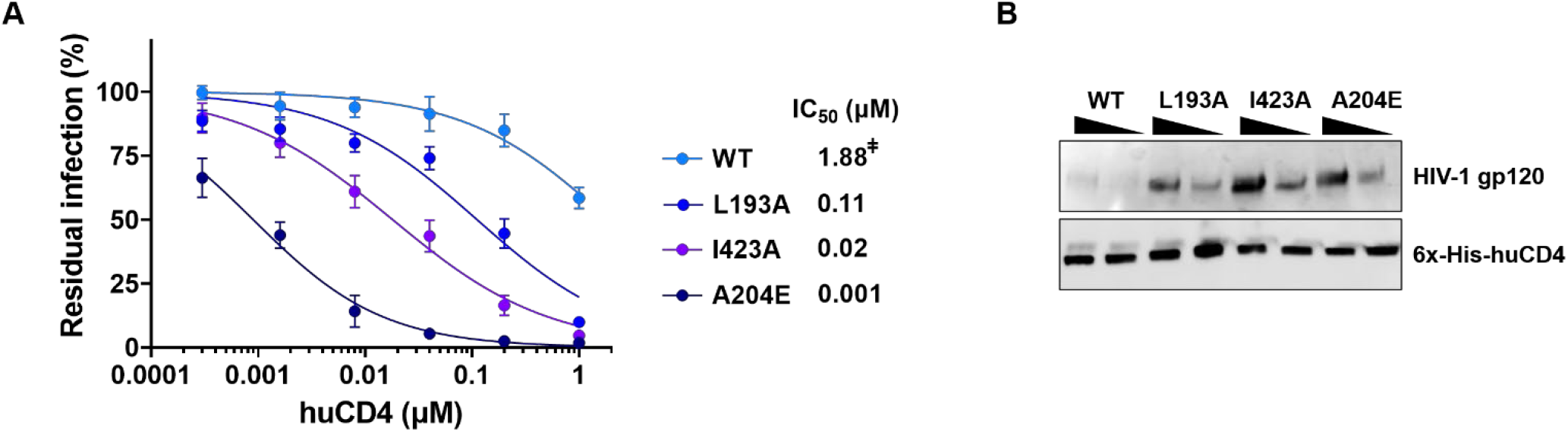
Effect of Env conformation on huCD4 binding. **(A)** Sensitivity of SHIV-BG505 to neutralization by soluble huCD4. Neutralization curves of the indicated SHIV variants were generated by plotting percent residual infection (y-axis) against huCD4 concentration (μM, x-axis). The key at the right of the graph indicates the identity of Env variant (wild-type ‘WT’ or indicated amino acid mutation) encoded by each SHIV. Each data point represents the average of two independent experiments, performed in duplicate. Error bars represent standard deviations. The calculated IC_50_s are shown. Where IC_50_ values were above the highest tested concentration, the extrapolated concentration is indicated by ^‡^. **(B)** Binding of SHIV-BG505 to huCD4. Western blot analysis for affinity pull-down of His-tagged huCD4 with increasing amounts (400 and 800 mU of RT) of indicated SHIV virions. Immunoblotting performed using anti-HIV-1 gp120 and anti-6x-His antibodies.

## Acknowledgments

We thank Julie Overbaugh for supporting the development of rhCD4_LOW_ and rhCD4_HIGH_ cells lines described here (NIDA DP1 DA039543), Ronald Desrosiers for providing the immortalized 221 T-lymphocytes, and Kyle Yamat for technical assistance. The following reagents were obtained through the NIH HIV Reagent Program, Division of AIDS, NIAID, NIH: TZM-bl cells (contributed by John Kappes and Xiaoyun Wu), Cf2Th/syn CCR5 cells (contributed by Tajib Mirzabekov and Joseph Sodroski), and HIV-1 gp120 antiserum (contributed by Michael Phelan).

## Author contributions

Conceptualization: AS; research design: AH, AS; investigation: GVV, ACS, HCB, AS; contributed methods/reagents: RKS, RCL; writing—original draft: AS; writing—review and editing: all authors.

## References

1. Hatziioannou T, Evans DT. 2012. Animal models for HIV/AIDS research. Nat Rev Microbiol 10:852–67.

2. Sharma A, Boyd DF, Overbaugh J. 2015. Development of SHIVs with circulating, transmitted HIV-1 variants. J Med Primatol 44:296–300.

3. Chen Z, Huang Y, Zhao X, Skulsky E, Lin D, Ip J, Gettie A, Ho DD. 2000. Enhanced infectivity of an R5-tropic simian/human immunodeficiency virus carrying human immunodeficiency virus type 1 subtype C envelope after serial passages in pig-tailed macaques (Macaca nemestrina). J Virol 74:6501–10.

4. Harouse JM, Gettie A, Eshetu T, Tan RC, Bohm R, Blanchard J, Baskin G, Cheng-Mayer C. 2001. Mucosal transmission and induction of simian AIDS by CCR5-specific simian/human immunodeficiency virus SHIV(SF162P3). J Virol 75:1990–5.

5. Ndung’u T, Lu Y, Renjifo B, Touzjian N, Kushner N, Pena-Cruz V, Novitsky VA, Lee TH, Essex M. 2001. Infectious simian/human immunodeficiency virus with human immunodeficiency virus type 1 subtype C from an African isolate: rhesus macaque model. J Virol 75:11417–25.

6. Nishimura Y, Shingai M, Willey R, Sadjadpour R, Lee WR, Brown CR, Brenchley JM, Buckler-White A, Petros R, Eckhaus M, Hoffman V, Igarashi T, Martin MA. 2010. Generation of the pathogenic R5-tropic simian/human immunodeficiency virus SHIVAD8 by serial passaging in rhesus macaques. J Virol 84:4769–81.

7. Pal R, Taylor B, Foulke JS, Woodward R, Merges M, Praschunus R, Gibson A, Reitz M. 2003. Characterization of a simian human immunodeficiency virus encoding the envelope gene from the CCR5-tropic HIV-1 Ba-L. J Acquir Immune Defic Syndr 33:300–7.

8. Song RJ, Chenine AL, Rasmussen RA, Ruprecht CR, Mirshahidi S, Grisson RD, Xu W, Whitney JB, Goins LM, Ong H, Li PL, Shai-Kobiler E, Wang T, McCann CM, Zhang H, Wood C, Kankasa C, Secor WE, McClure HM, Strobert E, Else JG, Ruprecht RM. 2006. Molecularly cloned SHIV-1157ipd3N4: a highly replication-competent, mucosally transmissible R5 simian-human immunodeficiency virus encoding HIV clade C Env. J Virol 80:8729–38.

9. Sharma A, Overbaugh J. 2018. CD4-HIV-1 Envelope Interactions: Critical Insights for the Simian/HIV/Macaque Model. AIDS Res Hum Retroviruses 34:778–779.

10. Boyd DF, Peterson D, Haggarty BS, Jordan AP, Hogan MJ, Goo L, Hoxie JA, Overbaugh J. 2015. Mutations in HIV-1 envelope that enhance entry with the macaque CD4 receptor alter antibody recognition by disrupting quaternary interactions within the trimer. J Virol 89:894–907.

11. Munro JB, Gorman J, Ma X, Zhou Z, Arthos J, Burton DR, Koff WC, Courter JR, Smith AB, 3rd, Kwong PD, Blanchard SC, Mothes W. 2014. Conformational dynamics of single HIV-1 envelope trimers on the surface of native virions. Science 346:759–63.

12. Ma X, Lu M, Gorman J, Terry DS, Hong X, Zhou Z, Zhao H, Altman RB, Arthos J, Blanchard SC, Kwong PD, Munro JB, Mothes W. 2018. HIV-1 Env trimer opens through an asymmetric intermediate in which individual protomers adopt distinct conformations. Elife 7.

13. Wang Q, Finzi A, Sodroski J. 2020. The Conformational States of the HIV-1 Envelope Glycoproteins. Trends Microbiol 28:655–667.

14. Kwon YD, Finzi A, Wu X, Dogo-Isonagie C, Lee LK, Moore LR, Schmidt SD, Stuckey J, Yang Y, Zhou T, Zhu J, Vicic DA, Debnath AK, Shapiro L, Bewley CA, Mascola JR, Sodroski JG, Kwong PD. 2012. Unliganded HIV-1 gp120 core structures assume the CD4-bound conformation with regulation by quaternary interactions and variable loops. Proc Natl Acad Sci U S A 109:5663–8.

15. Herschhorn A, Gu C, Moraca F, Ma X, Farrell M, Smith AB, 3rd, Pancera M, Kwong PD, Schon A, Freire E, Abrams C, Blanchard SC, Mothes W, Sodroski JG. 2017. The beta20-beta21 of gp120 is a regulatory switch for HIV-1 Env conformational transitions. Nat Commun 8:1049.

16. Herschhorn A, Ma X, Gu C, Ventura JD, Castillo-Menendez L, Melillo B, Terry DS, Smith AB, 3rd, Blanchard SC, Munro JB, Mothes W, Finzi A, Sodroski J. 2016. Release of gp120 Restraints Leads to an Entry-Competent Intermediate State of the HIV-1 Envelope Glycoproteins. mBio 7.

17. Humes D, Emery S, Laws E, Overbaugh J. 2012. A species-specific amino acid difference in the macaque CD4 receptor restricts replication by global circulating HIV-1 variants representing viruses from recent infection. J Virol 86:12472–83.

18. Keele BF, Giorgi EE, Salazar-Gonzalez JF, Decker JM, Pham KT, Salazar MG, Sun C, Grayson T, Wang S, Li H, Wei X, Jiang C, Kirchherr JL, Gao F, Anderson JA, Ping LH, Swanstrom R, Tomaras GD, Blattner WA, Goepfert PA, Kilby JM, Saag MS, Delwart EL, Busch MP, Cohen MS, Montefiori DC, Haynes BF, Gaschen B, Athreya GS, Lee HY, Wood N, Seoighe C, Perelson AS, Bhattacharya T, Korber BT, Hahn BH, Shaw GM. 2008. Identification and characterization of transmitted and early founder virus envelopes in primary HIV-1 infection. Proc Natl Acad Sci U S A 105:7552–7.

19. Han Q, Jones JA, Nicely NI, Reed RK, Shen X, Mansouri K, Louder M, Trama AM, Alam SM, Edwards RJ, Bonsignori M, Tomaras GD, Korber B, Montefiori DC, Mascola JR, Seaman MS, Haynes BF, Saunders KO. 2019. Difficult-to-neutralize global HIV-1 isolates are neutralized by antibodies targeting open envelope conformations. Nat Commun 10:2898.

20. Humes D, Overbaugh J. 2011. Adaptation of subtype a human immunodeficiency virus type 1 envelope to pig-tailed macaque cells. J Virol 85:4409–20.

21. Alexander L, Du Z, Rosenzweig M, Jung JU, Desrosiers RC. 1997. A role for natural simian immunodeficiency virus and human immunodeficiency virus type 1 nef alleles in lymphocyte activation. J Virol 71:6094–9.

22. Finzi A, Xiang SH, Pacheco B, Wang L, Haight J, Kassa A, Danek B, Pancera M, Kwong PD, Sodroski J. 2010. Topological layers in the HIV-1 gp120 inner domain regulate gp41 interaction and CD4-triggered conformational transitions. Mol Cell 37:656–67.

23. Haim H, Strack B, Kassa A, Madani N, Wang L, Courter JR, Princiotto A, McGee K, Pacheco B, Seaman MS, Smith AB, 3rd, Sodroski J. 2011. Contribution of intrinsic reactivity of the HIV-1 envelope glycoproteins to CD4-independent infection and global inhibitor sensitivity. PLoS Pathog 7:e1002101.

24. Kassa A, Madani N, Schon A, Haim H, Finzi A, Xiang SH, Wang L, Princiotto A, Pancera M, Courter J, Smith AB, 3rd, Freire E, Kwong PD, Sodroski J. 2009. Transitions to and from the CD4-bound conformation are modulated by a single-residue change in the human immunodeficiency virus type 1 gp120 inner domain. J Virol 83:8364–78.

25. Xiang SH, Kwong PD, Gupta R, Rizzuto CD, Casper DJ, Wyatt R, Wang L, Hendrickson WA, Doyle ML, Sodroski J. 2002. Mutagenic stabilization and/or disruption of a CD4-bound state reveals distinct conformations of the human immunodeficiency virus type 1 gp120 envelope glycoprotein. J Virol 76:9888–99.

26. Joag SV, Li Z, Foresman L, Stephens EB, Zhao LJ, Adany I, Pinson DM, McClure HM, Narayan O. 1996. Chimeric simian/human immunodeficiency virus that causes progressive loss of CD4+ T cells and AIDS in pig-tailed macaques. J Virol 70:3189–97.

27. Tan RC, Harouse JM, Gettie A, Cheng-Mayer C. 1999. In vivo adaptation of SHIV(SF162): chimeric virus expressing a NSI, CCR5-specific envelope protein. J Med Primatol 28:164–8.

28. Ren W, Mumbauer A, Gettie A, Seaman MS, Russell-Lodrigue K, Blanchard J, Westmoreland S, Cheng-Mayer C. 2013. Generation of lineage-related, mucosally transmissible subtype C R5 simian-human immunodeficiency viruses capable of AIDS development, induction of neurological disease, and coreceptor switching in rhesus macaques. J Virol 87:6137–49.

29. Li H, Wang S, Kong R, Ding W, Lee FH, Parker Z, Kim E, Learn GH, Hahn P, Policicchio B, Brocca-Cofano E, Deleage C, Hao X, Chuang GY, Gorman J, Gardner M, Lewis MG, Hatziioannou T, Santra S, Apetrei C, Pandrea I, Alam SM, Liao HX, Shen X, Tomaras GD, Farzan M, Chertova E, Keele BF, Estes JD, Lifson JD, Doms RW, Montefiori DC, Haynes BF, Sodroski JG, Kwong PD, Hahn BH, Shaw GM. 2016. Envelope residue 375 substitutions in simian-human immunodeficiency viruses enhance CD4 binding and replication in rhesus macaques. Proc Natl Acad Sci U S A 113:E3413–22.

30. Del Prete GQ, Ailers B, Moldt B, Keele BF, Estes JD, Rodriguez A, Sampias M, Oswald K, Fast R, Trubey CM, Chertova E, Smedley J, LaBranche CC, Montefiori DC, Burton DR, Shaw GM, Markowitz M, Piatak M, Jr., KewalRamani VN, Bieniasz PD, Lifson JD, Hatziioannou T. 2014. Selection of unadapted, pathogenic SHIVs encoding newly transmitted HIV-1 envelope proteins. Cell Host Microbe 16:412–8.

31. Chang HW, Tartaglia LJ, Whitney JB, Lim SY, Sanisetty S, Lavine CL, Seaman MS, Rademeyer C, Williamson C, Ellingson-Strouss K, Stamatatos L, Kublin J, Barouch DH. 2015. Generation and evaluation of clade C simian-human immunodeficiency virus challenge stocks. J Virol 89:1965–74.

32. Wei X, Decker JM, Liu H, Zhang Z, Arani RB, Kilby JM, Saag MS, Wu X, Shaw GM, Kappes JC. 2002. Emergence of resistant human immunodeficiency virus type 1 in patients receiving fusion inhibitor (T-20) monotherapy. Antimicrob Agents Chemother 46:1896–905.

33. Farzan M, Mirzabekov T, Kolchinsky P, Wyatt R, Cayabyab M, Gerard NP, Gerard C, Sodroski J, Choe H. 1999. Tyrosine sulfation of the amino terminus of CCR5 facilitates HIV-1 entry. Cell 96:667–76.

34. Nahabedian J, Sharma A, Kaczmarek ME, Wilkerson GK, Sawyer SL, Overbaugh J. 2017. Owl monkey CCR5 reveals synergism between CD4 and CCR5 in HIV-1 entry. Virology 512:180–186.

35. Sharma A, McLaughlin RN, Jr., Basom RS, Kikawa C, OhAinle M, Yount JS, Emerman M, Overbaugh J. 2019. Macaque interferon-induced transmembrane proteins limit replication of SHIV strains in an Envelope-dependent manner. PLoS Pathog 15:e1007925.

36. Meyerson NR, Sharma A, Wilkerson GK, Overbaugh J, Sawyer SL. 2015. Identification of Owl Monkey CD4 Receptors Broadly Compatible with Early-Stage HIV-1 Isolates. J Virol 89:8611–22.

37. Wu X, Parast AB, Richardson BA, Nduati R, John-Stewart G, Mbori-Ngacha D, Rainwater SM, Overbaugh J. 2006. Neutralization escape variants of human immunodeficiency virus type 1 are transmitted from mother to infant. J Virol 80:835–44.

38. Boyd DF, Sharma A, Humes D, Cheng-Mayer C, Overbaugh J. 2016. Adapting SHIVs In Vivo Selects for Envelope-Mediated Interferon-alpha Resistance. PLoS Pathog 12:e1005727.

39. Cavrois M, De Noronha C, Greene WC. 2002. A sensitive and specific enzyme-based assay detecting HIV-1 virion fusion in primary T lymphocytes. Nat Biotechnol 20:1151–4.

40. Roesch F, OhAinle M, Emerman M. 2018. A CRISPR screen for factors regulating SAMHD1 degradation identifies IFITMs as potent inhibitors of lentiviral particle delivery. Retrovirology 15:26.

41. Selig L, Pages JC, Tanchou V, Preveral S, Berlioz-Torrent C, Liu LX, Erdtmann L, Darlix J, Benarous R, Benichou S. 1999. Interaction with the p6 domain of the gag precursor mediates incorporation into virions of Vpr and Vpx proteins from primate lentiviruses. J Virol 73:592–600.

42. Vermeire J, Naessens E, Vanderstraeten H, Landi A, Iannucci V, Van Nuffel A, Taghon T, Pizzato M, Verhasselt B. 2012. Quantification of reverse transcriptase activity by real-time PCR as a fast and accurate method for titration of HIV, lenti- and retroviral vectors. PLoS One 7:e50859.

